# A mutation in *Themis* contributes to peanut-induced oral anaphylaxis in CC027 mice

**DOI:** 10.1101/2023.09.13.557467

**Authors:** Ellen L. Risemberg, Johanna M. Smeekens, Marta C. Cruz Cisneros, Brea K. Hampton, Pablo Hock, Colton L. Linnertz, Darla R. Miller, Kelly Orgel, Ginger D. Shaw, Fernando Pardo Manuel de Villena, A. Wesley Burks, William Valdar, Michael D. Kulis, Martin T. Ferris

## Abstract

**Background:** The development of peanut allergy is due to a combination of genetic and environmental factors, although specific genes have proven difficult to identify. Previously, we reported that peanut-sensitized CC027/GeniUnc (CC027) mice develop anaphylaxis upon oral challenge to peanut, unlike C3H/HeJ (C3H) mice.

**Objective:** To determine the genetic basis of orally-induced anaphylaxis to peanut in CC027 mice.

**Methods:** A genetic mapping population between CC027 and C3H mice was designed to identify the genetic factors that drive oral anaphylaxis. A total of 356 CC027xC3H backcrossed mice were generated, sensitized to peanut, then challenged to peanut by oral gavage. Anaphylaxis and peanut-specific IgE were quantified for all mice. T-cell phenotyping was conducted on CC027 and five additional CC strains.

**Results:** Anaphylaxis to peanut was absent in 77% of backcrossed mice, with 19% showing moderate anaphylaxis, and 4% having severe anaphylaxis. A total of eight genetic loci were associated with variation in response to peanut challenge, six associated with anaphylaxis (temperature decrease) and two associated with peanut-specific IgE levels. There were two major loci that impacted multiple aspects of the severity of acute anaphylaxis, at which the CC027 allele was associated with worse outcome. At one of these loci, CC027 has a private genetic variant in the *Themis* (thymocyte-expressed molecule involved in selection) gene. Consistent with *Themis*’ described functions, we found that CC027 have more immature T cells with fewer CD8+, CD4+, and CD4+CD25+CD127- regulatory T cells.

**Conclusion:** Our results demonstrate a key role for *Themis* in the orally-reactive CC027 mouse model of peanut allergy.

## INTRODUCTION

Food allergy is an immunologic disease with a significant public health and economic burden, affecting approximately 10% of adults^1^ and 8% of children^2,3^ in the US. Its prevalence is growing across the globe^4,5^, increasing the urgency of understanding its etiology. Peanut allergy is particularly severe as it is associated with high rates of anaphylaxis^1,6,7^, affects at least 1% of the US population^1,2,8^, and it is often lifelong, with 75-80% of pediatric cases persisting into adulthood^9–11^. Food allergy results from antigen-specific Th2 skewing leading to allergen-specific IgE production and is driven by complex interactions between host genetics and environmental factors^12^. Despite increased understanding of the environmental risk factors for food allergy development^13–15^, the genetic factors leading to these aberrant immune responses and the ways they interact with environmental factors are largely unknown.

Several studies have shown that immune homeostasis and the tendency for aberrant immune responses in a variety of diseases are under genetic control^16–21^. Similarly, studies in humans and mice have demonstrated that peanut allergy is highly heritable^22,23^ and the predisposition for individuals to develop peanut allergy is at least partially due to genetic variation^24,25^. Multiple studies have shown that genetic variation at the HLA locus^26–28^ and the filaggrin gene^29,30^ are associated with predisposition to peanut allergy in humans. However, these loci do not explain all the heritable variation in susceptibility to peanut allergy, nor in severity of allergic reaction. Atopic diseases in general are complex and polygenic, as evidenced by the finding of many genetic risk loci each with small effect^31^, and challenges remain in identifying additional associated genes.

Human studies on the genetics of food allergy to date are limited in their statistical power by sample sizes and the presence of confounding environmental variables. Highly powered, directed genetic studies are possible using experimental model organisms that recapitulate the development of peanut allergy in humans. Such model organisms also facilitate characterization of mechanisms underlying the development of peanut allergy and can be used to assess preclinical prophylactics and therapeutics. In particular, the laboratory mouse has become a widely used food allergy model that mimics key elements of the disease in humans^32^. In mouse models, sensitization and exposure can be tightly controlled, the genomes are well-characterized, and a wider range of tissues can be sampled (such as lymph nodes and spleen) to study the allergic response in more depth^33^. We have recently identified the Collaborative Cross^34,35^ strain CC027/GeniUnc (CC027) as an improved mouse model of peanut allergy^25^. Specifically, we showed that unlike previous mouse models, this strain develops anaphylaxis upon oral challenge and can be sensitized to peanut as an allergen in the absence of T_H_2-skewing adjuvants.

Here, we extend upon this discovery to dissect the genetic basis underlying CC027’s unique susceptibility to orally-induced peanut anaphylaxis. We performed a backcross between CC027, which reacts to oral peanut challenge, and C3H/HeJ (C3H), a long-standing mouse model for peanut allergy which requires challenge via intraperitoneal (i.p.) injection to elicit an allergic reaction. Using Quantitative Trait Locus (QTL) mapping, we find that there are several genetic loci associated with severity of anaphylaxis following oral challenge, with three major effect loci driving multiple aspects of anaphylaxis severity. One of these loci has a strong candidate mutation in the T-cell developmental gene *Themis*, and further characterization of CC027 demonstrates alterations in its T-cell compartments.

Altogether, our results point to a key immunologic mechanism for *Themis* in peanut allergy.

## METHODS

### Mice

CC027 mice were purchased from the Systems Genetics Core Facility at UNC Chapel Hill between 2015 and 2019. C3H mice were purchased from the Jackson Laboratory and bred at UNC for multiple generations between 2014 and 2019. CC027 and C3H colonies were maintained in the same animal facility at UNC. All mice were kept on a 12-hour light/dark cycle and had *ad libitum* access to peanut-free food and water. A backcross population was generated by first breeding CC027 and C3H mice together in both directions (i.e., C3H dam to CC027 sires, and CC027 dams to C3H sires). These F1 mice were then bred back to CC027 (again in both directions), such that the backcross population included animals with all four potential grandparental combinations. In total, we generated 356 female backcross mice for this study. At weaning, backcross mice were ear punched with a unique identifying number, had tail clips taken for DNA extraction and downstream genotyping, and were randomized across experimental cages to minimize the effects of litter or cross type on phenotypic outcomes. Between 4-6 weeks of age, mice were transferred from the breeding facility to another facility at UNC where they underwent the sensitization protocol (described below).

### Peanut Sensitization and Challenge

Peanut proteins were extracted from roasted defatted peanut flour (Golden Peanut, Alpharetta, GA) in PBS with 1 M NaCl, as previously described^36^. Female backcross mice, 4-6 weeks of age, were sensitized to peanut by oral gavage with peanut extract plus cholera toxin (List Biological Labs, Campbell, CA) once per week for four weeks. In the first three weeks, mice were given 2 mg peanut protein plus 10 µg cholera toxin, and in the fourth week, mice were given 5 mg peanut protein plus 10 µg cholera toxin.

One week following the final sensitization, mice were bled via the submandibular junction, and serum was isolated for subsequent immunoglobulin quantification. The following day, mice were challenged to peanut by oral gavage of 9 mg peanut protein in PBS. Anaphylaxis was monitored by measuring core body temperatures using a rectal thermometer (Physitemp, Clifton, NJ) every 15 minutes for one hour post-challenge. Symptom scores were also recorded as a readout of reaction severity using the following scale: 0, No symptoms; 1, Scratching and rubbing around the nose and head; 2, Puffiness around eyes and mouth, diarrhea, pilar erecti, reduced activity, and/or decreased activity with increased respiratory rate; 3, Wheezing, labored respiration, cyanosis around mouth, feet, tail; 4, No activity after prodding; tremor or convulsion; 5, Death.

### Genotyping

DNA was extracted from tail biopsies using Qiagen (Qiagen, Inc) DNeasy extraction. 500 ng of DNA were submitted to Neogen (Lincoln, NE) for running on the MiniMUGA genotyping array^37^. We filtered the 10,819 biallelic single nucleotide polymorphisms (SNPs) to an informative and well-performing set of 2,890 based on the following criteria: SNPs 1) segregated between the parental CC027 and C3H strains, 2) had an ‘N’ call rate of less than 5%, 3) had close to expected genotype frequencies of 50% CC027/CC027 and 50% CC027/C3H (between 30% and 70% homozygous).

### T-cell Analysis

Male and female mice from five CC strains (CC007, CC019, CC026, CC027, CC061) between 4-5 weeks of age were ordered from the UNC Systems Genetics Core Facility in 2022. Mice were kept on a 12:12 light/dark cycle with *ad libitum* access to peanut-free food and water. At 6-7 weeks of age, mice were euthanized via isoflurane overdose. Thymus, mesenteric lymph nodes, and spleens were isolated and placed in cold RPMI + 1% FBS and kept on ice. Cells were plated from single cell suspensions in FACS buffer (HBSS+2% FBS) in a 96-well round bottom plate. Cells were incubated for 45 minutes at 4C for antibody staining with the following antibodies: Live/Dead Fixable Aqua, CD3-APC/Cy7 (145-2C11), CD127-PerCP/eF710 (eBioSB/199), CD4-PE (GK1.5), CD8-BV650 (53-6.7), and CD25-BV711 (PC61).

Following staining, cells were washed twice with HBSS, fixed with 2% PFA (in PBS), and stored at 4°C in the dark until analyzed on a Beckman Coulter Attune NxT flow cytometer. Resultant data were analyzed in FlowJo to assess the overall abundance and relative proportions of multiple T-cell populations.

### Data Analysis

#### Heritability

Heritability for each phenotype was estimated in two ways: 1) by calculating the proportion of phenotypic variation explained by strain in inbred parents (ANOVA) and 2) by estimating the proportion of phenotypic variance explained by the additive effects at all genotyped SNPs (additive SNP heritability^38^) using a linear mixed model with a polygenic random effect. The genomic relationship matrix of the random effect was defined using GCTA-GRM^39^ and the lme4qtl R package^40^ was used to estimate the associated variance component.

#### Quantitative Trait Loci (QTL) Mapping

We performed QTL mapping using Haley-Knott regression^41^ with the R/qtl R package^42^. Cage and batch were included as random effect covariates and baseline temperature was included as a fixed effect covariate for temperature-related phenotypes. Significance thresholds were determined by permutation test, with maximum LOD scores fit to a Generalized Extreme Value (GEV) distribution^43,44^ using the extRemes R package^45^. Bayesian credible intervals were calculated for QTLs as described by Sen & Churchill (2001)^46^.

#### Candidate Gene Analysis

Following identification of QTL, relevant genetic and genomic data were used to identify candidate genes driving the association between QTL and phenotype. First, we identified all single nucleotide polymorphisms (SNPs) that varied between C3H and the CC027 founder strain haplotype(s) present at the locus^47^ and annotated them with SnpEff^48^. Variants were categorized as protein-modifying or regulatory based on sequence ontology descriptions^49^ and counts were obtained of genes containing these protein-modifying or regulatory variants. Variant data was obtained from the Mouse Genomes Project (MGP)^50^. GATK SelectVariants tool (v3.7)^51^ was used to select variants between the two strains of interest for each QTL region, where variants were biallelic snps or indels. Ensembl IDs were mapped to gene names using the GRCm38 build (release 79) of *mus musculus*.

## RESULTS

### CC027xC3H backcross mice show highly variable and heritable responses to oral peanut challenge

Our lab previously showed that the temperature trajectories of CC027 and C3H mice are significantly different^25^ (**Fig. 1A**, p=0.004, repeated measures ANOVA). Here we generated a CC027xC3H backcross population of 356 female mice and evaluated their responses following sensitization and oral peanut challenge. Temperature was measured every 15 minutes for one hour post-challenge, since a major symptom of anaphylaxis in mice is hypothermia. We calculated an “area above the curve” (AAC) summary statistic representing the overall temperature trajectory (see Supplemental Methods and illustration in **Supp. Fig. 1**). This metric showed more variation in backcross mice than observed in the parent strains, as expected with a multi- or polygenic trait (**Fig. 1B**; p=0.07, Browne-Forsythe test).

**Figure 1.**
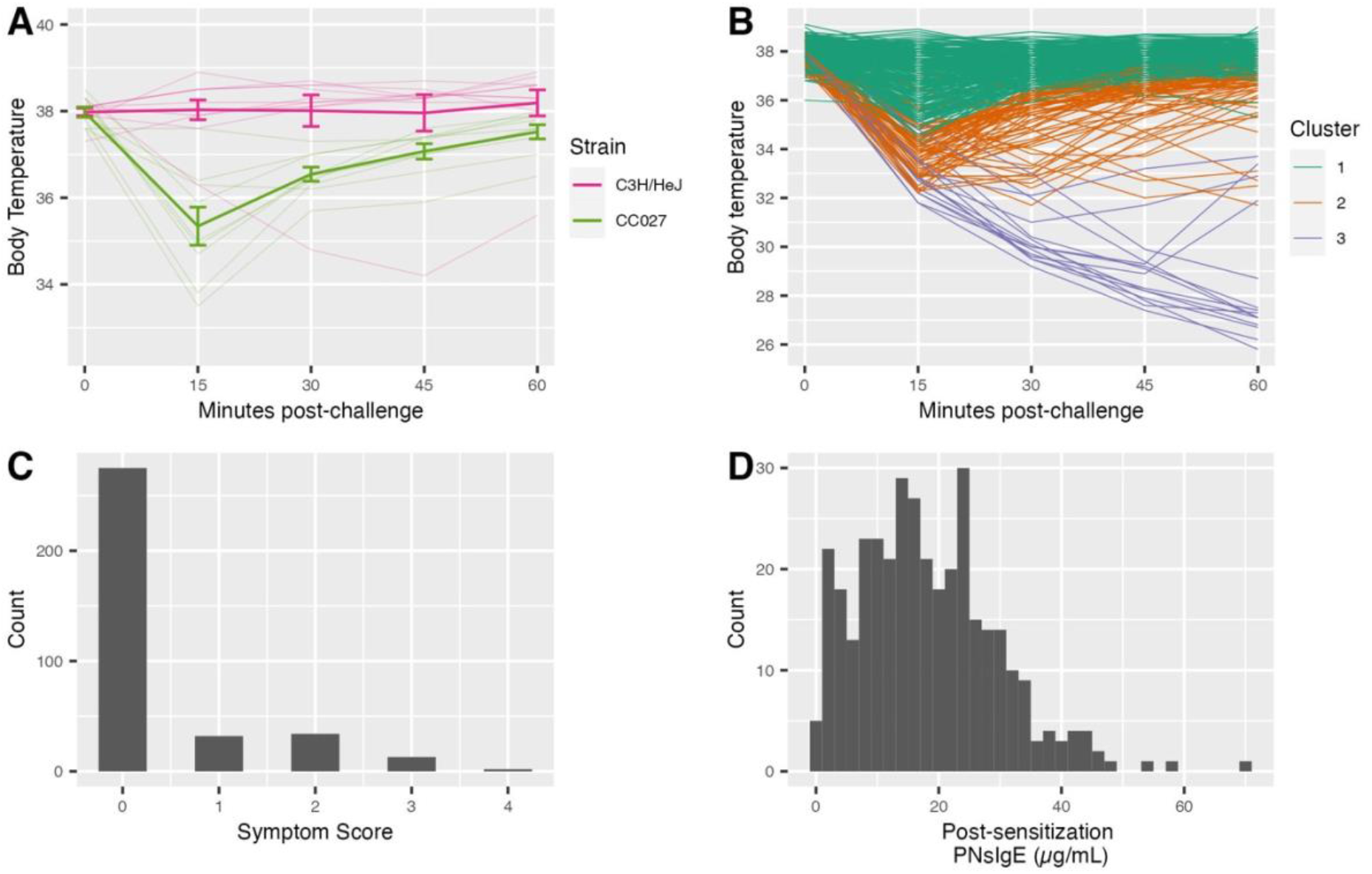
Food allergy-related phenotypes in C3H, CC027, and backcross mice. **A**) Temperature trajectory in parental strains (strain effect on trajectory: p = 0.004, repeated measures ANOVA). **B**) Temperature trajectory in backcross mice, colored by cluster from hierarchical clustering analysis based on temperature data. **C**) Symptom score distribution in backcross mice. **D**) Post-sensitization peanut-specific IgE (PNsIgE) distribution in backcross mice.

Hierarchical clustering based on observed temperature measurements divided these trajectories into three groups, with 77% (275) of the mice in the non-reactor (green) cluster, 19% (66) in the initial decrease and recovery (orange) cluster, and 4% (15) in the decrease and continued decline (purple) cluster (**Fig. 1B**).

The severity of allergy symptoms in the backcross mice was also measured with a 5-point symptom scale (see Methods) and found that 23% of the mice developed symptoms following challenge (**Fig. 1C**).

Lastly, to confirm all backcross mice had been sensitized to peanut, we measured serum peanut-specific IgE (PNsIgE) the day prior to challenge. All but two mice had detectable PNsIgE, and among mice with detectable PNsIgE, levels ranged from 0.7-70 μg/mL (**Fig. 1D**). Estimation of heritability showed that all allergic phenotypes were heritable, with additive heritability ranging from 0.18-0.76 in the inbred parents and 0.13-0.25 in the (backcross) offspring (**Supp. Table 1**).

Given the phenotypic ranges described, we sought to assess the relationships between phenotypes. The two measures of anaphylaxis, temperature trajectory and symptom score, were significantly associated (p<2×10^−6^, ANOVA) (**Supp. Fig. 2A**). Temperature trajectory was significantly but weakly associated with PNsIgE (Pearson’s r=0.16, p=0.002) (**Supp. Fig. 2B**). Together these observations and heritability calculations in the backcross mice support a wide range of responses to peanut challenge, driven in part by a genetic component.

### Susceptibility to orally-induced peanut anaphylaxis in CC027 mice is multigenic

Given the strong heritability and the range of phenotypes observed in the backcross, we sought to identify genetic loci associated with this phenotypic variation. Following genotyping of each backcross mouse, we filtered these genotypes down to a set of 2889 informative and reliable markers (see methods). These markers were well distributed across the genome (**Supp. Fig. 3**) giving us robust coverage to conduct genetic mapping. We performed QTL mapping on each of the observed phenotypes and summary statistics. Ten phenotypes were analyzed in total: temperature at 15, 30, 45, and 60 minutes; area above the curve (AAC) summarizing temperature trajectory; temperature trajectory cluster (binarized to non-reactor vs reactor); minimum temperature reached; time of minimum temperature reached; symptom score; and PNsIgE. We found eight genome-wide significant loci, each associated with at least one allergic phenotype, and named these *Qpa* (*QTL for peanut allergy*) *1-8*. QTL effects (see Supplemental Methods) accounted for between 3-8% of the total phenotypic variation observed in our backcross population (**Table 1**).

**Table 1.** Quantitative trait loci identified from mapping in backcross mice. Confidence intervals, genome-wide adjusted p-values, and proportion of phenotypic variance explained were calculated as described in methods and supplemental methods. In adjusted p-value column, * represents genome-wide adjusted p-value < 0.10, ** < 0.05, and *** < 0.005.

| QTL IDENTIFIER | ASSOCIATED PHENOTYPE(S) | CHR: POSITION (CONFIDENCE INTERVAL) (Mb) | ADJUSTED P-VALUE | PHENOTYPIC VARIATION EXPLAINED |
| --- | --- | --- | --- | --- |
| *Qpa1* | Temperature at 15 min | Chr 10: 28.39 (3.79-29.23) | 0.0032 (***) | 5.58% |
|  | Temperature at 30 min | Chr 10: 9.21 (3.79-29.23) | 0.015 (**) | 4.69% |
|  | Temperature - area under the curve (AAC) | Chr 10: 5.65 (3.79-29.23) | 0.025 (**) | 4.32% |
|  | Minimum temperature | Chr 10: 28.39 (3.79-29.23) | 0.028 (**) | 4.48% |
| *Qpa2* | Temperature trajectory cluster (non-reactors vs reactors) | Chr 5: 62.00 (26.75-93.05) | 0.021 (**) | 4.34% |
|  | Temperature at 45 min | Chr 5: 89.88 (16.84-106.93) | 0.036 (**) | 4.22% |
|  | Minimum temperature | Chr 5: 64.29 (15.88-112.94) | 0.059 (*) | 4.09% |
|  | Temperature - area under the curve (AAC) | Chr 5: 96.94 (20.80-114.72) | 0.084 (*) | 3.55% |
| *Qpa3* | Temperature at 30 min | Chr 17: 32.81 (17.85-49.48) | 0.0091 (**) | 4.48% |
|  | Minimum temperature | Chr 17: 32.81 (16.99-49.48) | 0.029 (**) | 4.12% |
|  | Temperature at 45 min | Chr 17: 35.55 (16.37-49.48) | 0.069 (*) | 3.47% |
|  | Symptom score | Chr 17: 35.55 (7.14-45.51) | 0.098 (*) | 4.26% |
| *Qpa4* | Time of minimum temperature | Chr 6: 129.82 (57.26-136.08) | 0.029 (**) | 3.74% |
| *Qpa5* | Minimum temperature | Chr 18: 89.55 (63.37-89.55) | 0.068 (*) | 3.68% |
| *Qpa6* | Temperature at 45 min | Chr 20: 108.94 (40.29-142.02) | 0.078 (*) | 3.06% |
| *Qpa7* | Post-sensitization peanut-specific IgE (µg/mL) | Chr 9: 95.33 (91.59-103.49) | 2.6e-05 (***) | 7.66% |
| *Qpa8* | Post-sensitization peanut-specific IgE (µg/mL) | Chr 11: 45.13 (37.02-69.76) | 0.0076 (**) | 4.79% |

Three of these loci, *Qpa1, Qpa2*, and *Qpa3* were defined as major effect loci due to their association with several allergy outcome phenotypes (**Table 1**). *Qpa1* (chr10:3.79-29.23 Mb) is most significantly associated with the initial temperature drop, *Qpa3* (chr5:26.75-93.05 Mb) with temperature at 30 minutes, and *Qpa2* (chr17:17.85-49.48 Mb) with temperature trajectory cluster (**Fig. 2A**). The allele effects of *Qpa1* and *Qpa2* are as expected with CC027 homozygotes having more severe hypothermia. These effects are reversed at *Qpa3*, with CC027 homozygotes having less severe hypothermia (**Fig 2**). Together these loci explained a significant amount of phenotypic variation present in the population, with the overall trajectory (represented by AAC) having 12.1% of its variation explained by these three loci. Remaining temperature-associated QTL (*Qpa4-6*) are summarized in **Supplemental Figure 4**.

**Figure 2.**
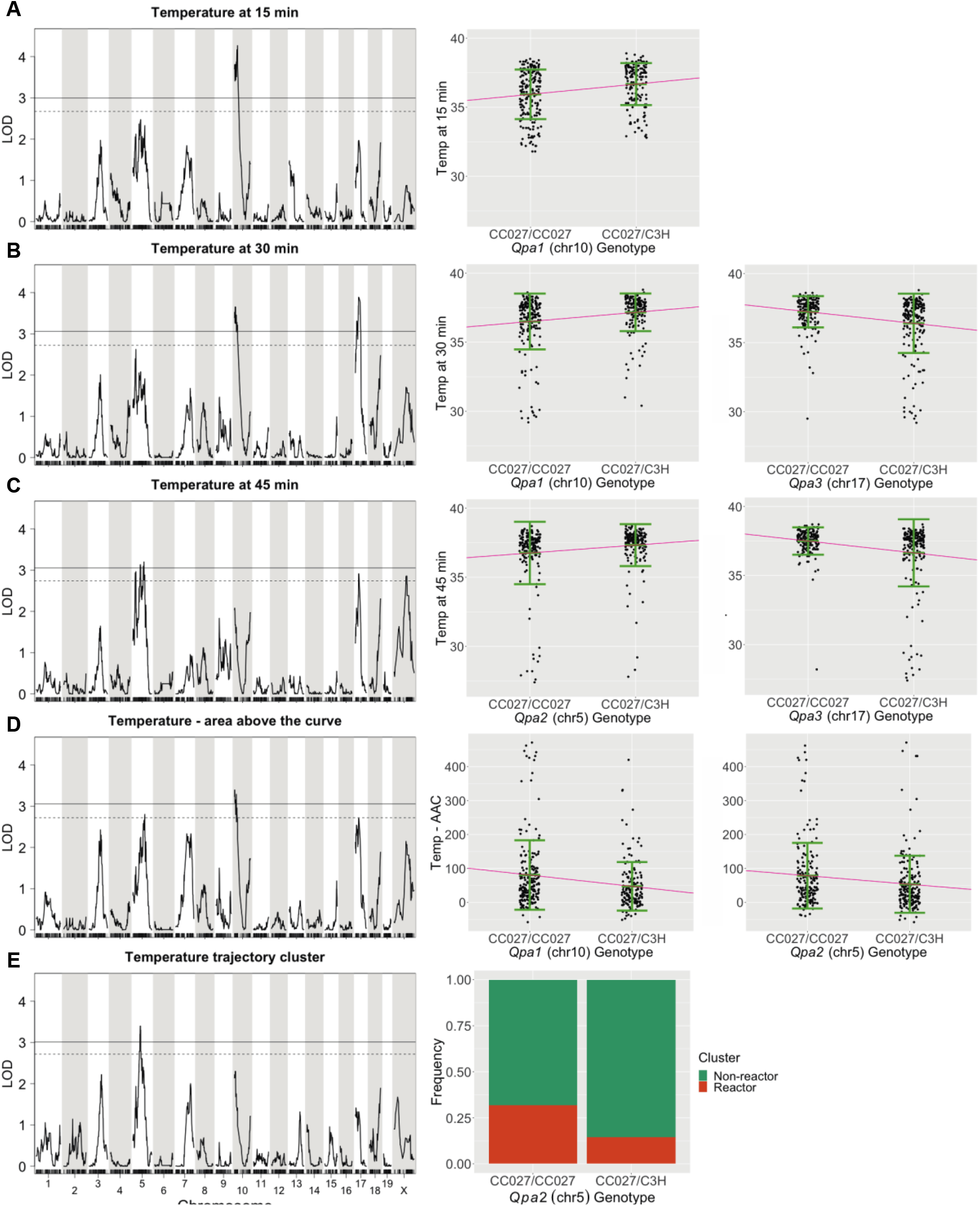
Genetic architecture of post-challenge temperature trajectory. Genome-wide scan for QTL associated with selected temperature phenotypes. Solid and dashed lines represent 5% and 10% significance thresholds, respectively. The right panel shows phenotype-by-genotype plots showing relationship between the genotype at each QTL and its associated phenotype. A) Temperature at 15 minutes. B) Temperature at 30 minutes. C) Temperature at 45 minutes. D) Temperature area above the curve. E) Temperature trajectory cluster (non-reactors vs reactors).

Given the number of loci identified, we sought to confirm the independence of each QTL’s effect on our trait(s) of interest. We first analyzed the cumulative effects of *Qpa1* and *Qpa2* (together explaining 8.6% of variation in AAC) based on the fact that they have consistent allele effects, with CC027 homozygotes at each locus having worse outcomes. Mice that were heterozygous at both loci had the least severe hypothermic response, and each additional CC027 allele at either locus resulted in incrementally increasing severity such that mice homozygous for CC027 at both loci had the most severe response (**Fig. 3**). The total number of CC027 alleles at *Qpa1* and *Qpa2* had a highly significant linear effect on AAC (p = 2.5×10^−6^, ANOVA) and there was no significant interaction between *Qpa1* and *Qpa2* (p = 0.94, ANOVA). As a complementary approach, we performed variable selection with stepwise regression to determine whether each QTL had an independently significant effect on severity of allergic reaction.

**Figure 3.**
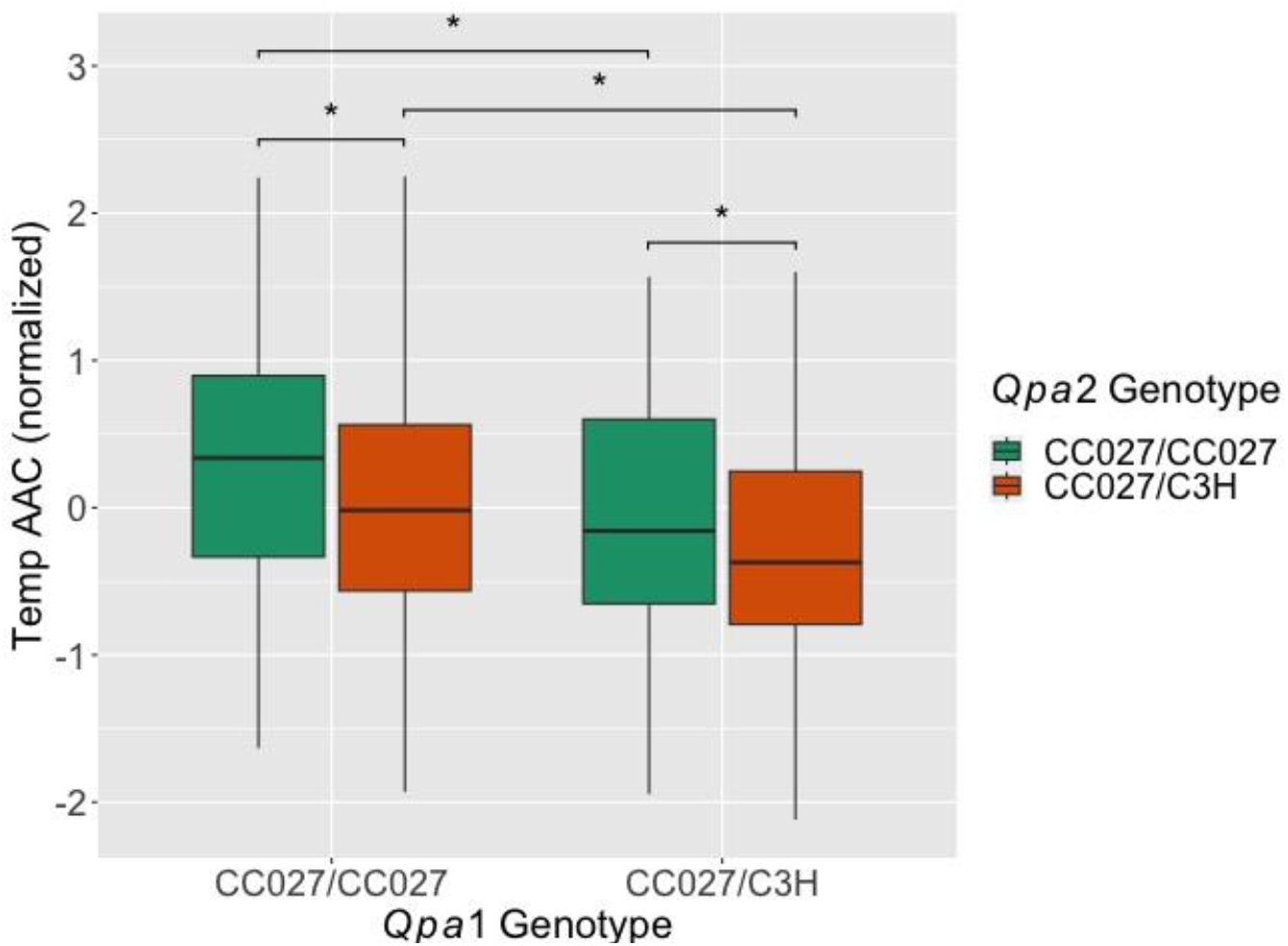
Cumulative additive effect of *Qpa1* and *Qpa2*. Boxplot showing AAC (rank inverse normalized so it is centered around zero) by genotype at *Qpa1* (x-axis) and *Qpa2* (color). Effect is additive with no significant interaction (p = 0.94, ANOVA). Mice that are homozygous CC027 at both loci (leftmost group) have a significantly higher AAC (larger temperature drop) than mice that are heterozygous at either locus (p = 0.02, 0.01; Welch’s two-sample t-test). AAC for mice that are heterozygous at either *Qpa1* or *Qpa2* (middle two groups) are not significantly different from each other (p = 0.81) and are significantly higher than AAC for mice that are heterozygous at both loci (rightmost group; p = 0.01, 0.03).

Using the AAC statistic as a representative outcome phenotype, we found that five of the six QTL associated with temperature, namely *Qpa1-6* except for *Qpa4*, were independently significant contributors to the linear model predicting AAC. PCA and factor analysis confirmed that *Qpa4*’s associated phenotype, time at minimum temperature, captures variation distinct from all other phenotypes (**Supp. Fig. 5**), thus explaining the exclusion of *Qpa4* from the model predicting AAC.

Lastly, we turned to our two loci affecting PNsIgE levels, *Qpa7* and *Qpa8* (**Fig 4**). At both loci, CC027 homozygotes have significantly less PNsIgE than heterozygotes. Together, these two loci explain 12% of the variation in PNsIgE. We also asked whether these loci affect temperature trajectory by assessing whether they improved the model predicting AAC. Although the PNsIgE phenotype itself improved the model (p=0.007, ANOVA), neither *Qpa7* (p=0.99, ANOVA) nor *Qpa8* (p=0.36, ANOVA) significantly improved the model predicting AAC.

**Figure 4.**
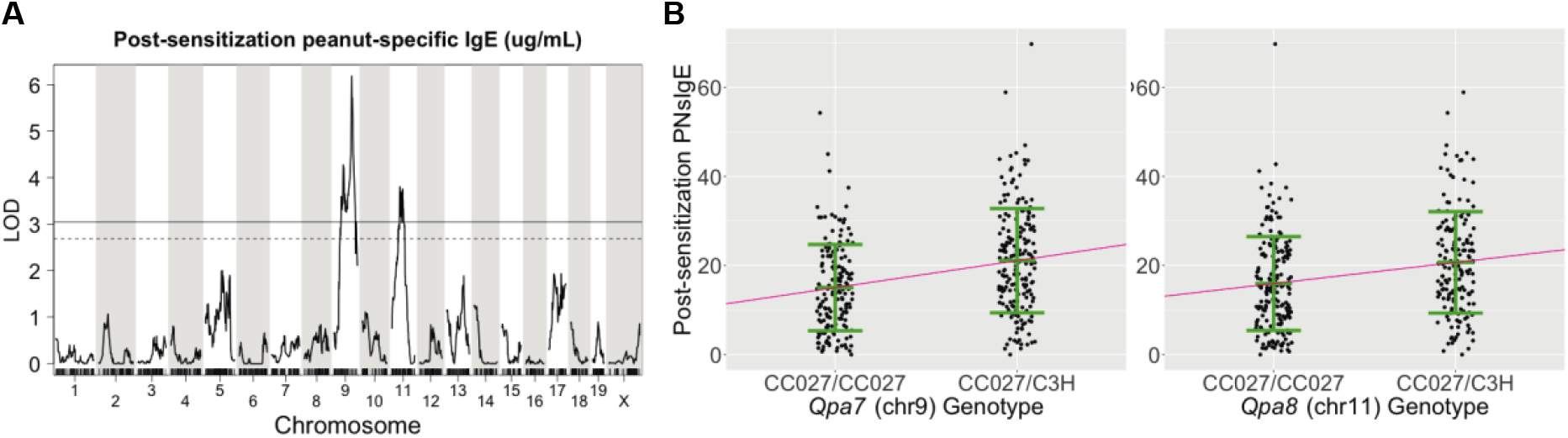
Genetic architecture of post-sensitization peanut-specific IgE. A) Genome-wide scan for QTL associated with post-sensitization peanut-specific IgE (PNsIgE). Solid and dashed lines represent 5% and 10% significance thresholds, respectively. B) Phenotype-by-genotype plots showing relationship between the genotype at each QTL and PNsIgE levels.

### CC027 harbors a T-cell compartment modifying mutation in *Themis*

To identify genes driving association between a locus and a phenotype, we looked for protein-modifying or regulatory variants segregating between C3H and CC027. We counted variants of each type as well as the associated genes within each QTL’s 95% credible interval (**Table 2**). The number of genes within each QTL was reduced by 20% on average, ranging from 0-82% reduction, when considering only those genes that contained such variants (compared with all genes in the region). Despite this narrowing of the candidate regions, there were at least 40 and up to 1162 candidate genes per locus. Notably, *Qpa7* contains 40 candidate genes, only 4 of which had protein-affecting variants, although none of these have obvious links to allergy development.

**Table 2.** Candidate gene analysis results. Region is the widest 95% confidence interval (CI) from all phenotypes associated with a QTL. Total Genes refers to the number of genes present in the 95% CI and Candidate Genes is the number of genes present after filtering for genes containing variants segregating between CC027 and C3H.

| QTL IDENTIFIER | CHR: POSITION | TOTAL GENES | CANDIDATE GENES | TOTAL VARIANTS (CC027 VS. C3H) | NUMBER OF CANDIDATE GENES |  |
| --- | --- | --- | --- | --- | --- | --- |
|  |  |  |  |  | WITH PROTEIN-CODING VARIANTS | WITH ONLY REGULATORY VARIANTS |
| *Qpa1* | Chr 10: 3.79-29.23 Mb | 258 | 147 | 19017 | 38 | 109 |
| *Qpa2* | Chr 5: 15.88-114.72 Mb | 1179 | 1141 | 287955 | 456 | 685 |
| *Qpa3* | Chr 17: 17.85-45.51 Mb | 979 | 975 | 70310 | 401 | 574 |
| *Qpa4* | Chr X: 40.29-142.02 Mb | 1553 | 861 | 54367 | 114 | 747 |
| *Qpa5* | Chr 6: 57.26-136.08 Mb | 1187 | 1162 | 226803 | 494 | 668 |
| *Qpa6* | Chr 18: 63.37-89.55 Mb | 221 | 220 | 103299 | 97 | 123 |
| *Qpa7* | Chr 9: 91.59-103.49 Mb | 217 | 40 | 1211 | 4 | 36 |
| *Qpa8* | Chr 11: 37.02-69.76 Mb | 759 | 715 | 60503 | 174 | 541 |

For most of the genomic region covered by the *Qpa1* locus (chr 10: 3.79-23 Mb), CC027 inherited a C57BL/6J (B6) haplotype, notable because B6 mice do not react to oral peanut challenge^25^. Given this result, we asked whether CC027 has any private (strain-specific) genetic variants within this region. Our lab previously reported that CC027 possesses a private variant intronic to the *Themis* gene in this region^52^. Given *Themis*’s known role in T-cell development^53–59^, we hypothesized that CC027 would exhibit aberrant T-cell development and would therefore vary in its T-cell compartment from strains without the *Themis* mutation. To study this, we measured the overall abundance and relative proportion of several T-cell subtypes (live, CD3+, CD4+, regulatory, CD8+, double negative and double positive) in the thymus, lymph nodes and spleen of 6-8 mice each from each of five CC strains: CC027, and four strains with the same (B6) haplotype as CC027 but not harboring the private mutation in *Themis* (CC007, CC019, CC026, and CC061).

Overall, CC027 had significantly more immature T cells and fewer mature T cells in the spleen, as well as more mature T cells in the thymus (**Fig. 5**). We first asked whether CC027 varied from the phenotypic average of the other strains using linear regression and found that CC027 was significantly different in all seven of the T-cell subtypes in the spleen, one subtype in the lymph nodes and three in the thymus (see Supplemental Methods, **Supp. Table 2**). In a more conservative test of pairwise comparisons (Tukey’s HSD^60^), CC027 varied from all four other strains in splenic Tregs (lower, **Fig. 5A**) and splenic double negative T cells (DNTs) (higher, **Fig. 5B**), and varied from three of the four strains in splenic CD8+ T-cells (lower, **Fig. 5C**) and thymic CD4+ T-cells (higher, Fig. 5D) (**Supp. Fig. 6, Supp. Table 2**). Altogether, these results suggest that CC027 harbors a variant T cell compartment compared to other strains with the same genetic background at *Qpa1* but without the private variant in *Themis*.

**Figure 5.**
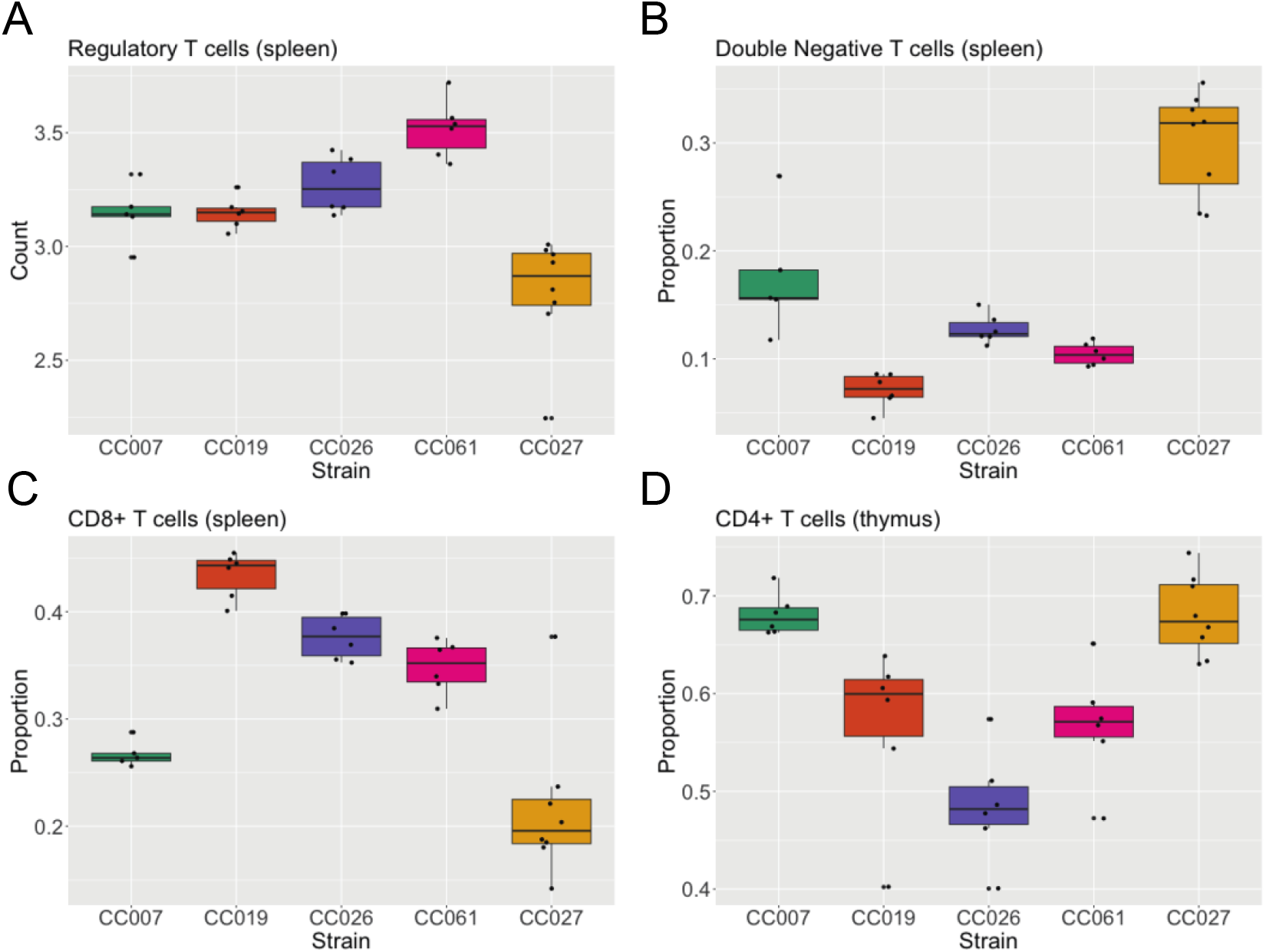
CC027 exhibits a modified T-cell compartment compared with strains not harboring the *Themis* mutation. Boxplots showing T-cell subtype measurements across five strains measured. A) Regulatory T cells in the spleen. B) double negative T cells in the spleen. C) CD8+ T cells in the spleen. D) CD4+ T cells in the spleen.

## DISCUSSION

A variety of environmental and genetic factors contribute to the likelihood of developing peanut allergy and determining the severity of allergic reaction^22–30^. Here we utilized CC027, our established mouse model of peanut allergy^25^, to dissect the genetic architecture underlying their susceptibility to anaphylaxis upon oral challenge. Our results show that the genetic component of this susceptibility is multigenic, with our study identifying eight loci significantly associated with various aspects of peanut-allergic responses. We identify *Themis* as a candidate gene underlying one of these associations, and in doing so identify a potential role for aberrant genetic regulation of T-cell development being a key driver of anaphylaxis severity in this model.

We identified a locus on chromosome 10, *Qpa1*, associated with multiple aspects of the early response to oral peanut challenge. *Qpa1* was most significantly associated with the initial response to oral challenge (temperature at 15 minutes post-challenge), suggesting *Qpa1* acts in the initial triggering of CC027’s orally-driven anaphylactic response to peanut. Although there are many genetic variants segregating between C3H and CC027 in this large (∼25 Mb) region, we were able to identify the likely candidate gene *Themis* (Thymocyte-Expressed Molecule Involved in Selection). CC027 has inherited CAST/EiJ (CAST) and B6 haplotypes in the *Qpa1* locus. Notably, other CC strains which inherited either CAST or B6 haplotype across this entire region were not susceptible to oral reactivity to peanut^25^, suggesting that these haplotypes alone are not sufficient to confer susceptibility to orally-driven allergy and the most likely cause is a *de novo* variant in the region private to CC027. *Themis* contains a private intronic variant in CC027^52^ and has a biological role in T-cell development^58,59,61^, which is relevant to the development of food allergy^62^. Further, variation in *Themis* has been associated with inflammatory disease in humans, including celiac disease^63,64^ and multiple sclerosis^65^, providing evidence for a role in immune-mediated human disease.

*Themis* has been shown specifically to be involved in promoting positive selection: the maturation of single positive T (SPT) cells from double positive T (DPT) cells before migration out of the thymus^53–58^. We therefore hypothesized that, relative to other strains containing a B6 haplotype at *Themis*, CC027 would likely show defects in some aspects of their T-cell compartment. Indeed, we found that CC027 was an outlier relative to these other strains in many aspects of their T-cell compartment. The most striking results, where CC027 varied significantly from all other strains, were that CC027 mice had both a significantly greater proportion of DNT cells and significantly fewer Tregs in the spleen. DNT cells are immature and should generally not be exported to peripheral lymphoid sites, so a higher proportion of DNT cells suggests aberrant T-cell development in the thymus. Tregs are a specific lineage of mature T cells that play a major role in immune homeostasis and the maintenance of self-tolerance^66^, notably in maintaining tolerance to potential allergens^67^. Decreased Tregs could play a direct role in the susceptibility of these mice to allergy. Furthermore, Tregs can directly inhibit mast cell degranulation through OX40-OX40L interactions^68^, and depleting Tregs leads to more severe anaphylaxis^68^, suggesting a direct link between the lower Treg quantities in CC027 mice and higher reactivity during oral peanut challenge. CC027 mice also have significantly fewer mature CD8+ and CD4+ T cells in the spleen and higher CD4+ T cells in the thymus, further suggesting aberrant T-cell development in the thymus. Together, these data support that *Themis* is a plausible causal gene for the effect of *Qpa1*.

We note that in these analyses, CC027 was more like CC007 than other strains in multiple aspects of their T-cell compartment. We have previously shown that there is a high degree of variation across the CC in different aspects of their T-cell compartment^20,69^ and their immune systems more generally^70–72^, providing a potential explanation for CC007’s apparent variant T-cell compartment. We did not formally test all the CC strains (including CC007) in our peanut allergy screen, so future work studying T-cell variation across the CC and how it relates to food allergy susceptibility will help characterizing genetic mechanisms of susceptibility. Future work will focus on formally studying the mechanism by which this *Themis* variant in CC027 is impacting the T-cell compartment, likely by examining expression in key T-cell populations during development and in response to challenge.

In addition to *Qpa1*, we identified five other loci associated with severity of anaphylactic response. These findings are in line with the polygenic nature of genetic contributions to food allergy in humans^12^. Similarly, the extreme response CC027 shows is likely unique because of the accumulation of multiple variants affecting allergic reaction severity. Because a backcross limits the number of recombinations to one chromosome (the one received from the F_1_ parent), QTL mapping in these populations can produce large QTL that are difficult to fine-map. Indeed, the QTL identified in this study were relatively large, making identification of specific candidates challenging except in the case of *Qpa1*. Therefore, future work will be focused on each of these loci individually (ideally in the context of a fixed *Themis* variant) to better characterize the genetic features and intermediate mechanisms that drive these associations with disease severity.

*Qpa3* is quite intriguing based on the finding that at this locus the heterozygous (C3H/CC027) animals were more susceptible than the CC027 homozygotes. This locus overlaps with the MHC region in the mouse genome (the equivalent of the human HLA region), which has been implicated in the development of peanut allergy in humans, so it is validating that we identified an association at this locus, albeit in an unexpected direction. The unexpected allele effects at this locus bring up the interesting possibility that many mouse strains may harbor genetic variants or loci that can contribute to susceptibility to allergy if they are placed in an otherwise permissive genomic contexts (in this case, having the *Qpa1* locus).

Several studies in humans^26–28^ have shown the importance of HLA haplotypes in predicting the likelihood of developing peanut allergy, highlighting the potential that adaptive immune recognition is a critical component of peanut allergy development. CC027, C3H and B6 all produce similar levels of PNsIgE^25^ despite their highly variant response to oral challenge. Though it was not surprising that we identified genetic loci affecting IgE levels in this cross, it was surprising that at these two loci, *Qpa7* and *Qpa8*, a heterozygous genotype (CC027/C3H) is associated with higher levels of PNsIgE relative to the CC027 homozygotes. Given the anticipated role of allergen-specific IgE in driving allergy development, it is somewhat surprising that CC027 homozygotes have less PNsIgE. However, prior research in our lab suggests that susceptibility to peanut allergy in mice is not explained by an increase in PNsIgE levels^25^, so our observations are not necessarily contradictory. We note that in our backcross population, PNsIgE is weakly (r=0.16) but significantly (p=0.002) correlated with severity of allergic response (as measured by AAC). Given this weak correlation along with our previous report that PNsIgE levels do not correlate with reaction severity in CC027 and other inbred strains^25^, differences beyond allergen-specific IgE levels (such as aberrant T-cell development) must be important for the severity of allergic reaction observed.

In summary, our results provide evidence for multigenic regulation of susceptibility to peanut allergy. We identify several loci associated with both severity of allergic response and peanut-specific IgE. We present a convincing causal gene in *Themis*, which may be affecting disease risk through aberrant regulation of T-cell development. This work identifying several loci and a specific candidate gene associated with peanut allergy follows the identification of CC027 as a model for peanut allergy, showing the importance of genetically diverse panels like the Collaborative Cross in having appropriate disease models. These results can be used to further understand the etiology of peanut allergy in humans and provides potential therapeutic targets in genes that regulate T-cell development.

## Abbreviations used

IP: intraperitoneal
C3H: C3H/HeJ
CC027: CC027/GeniUnc
QTL: quantitative trait locus

## DATA AVAILABILITY

All data that support the findings of this study and code to reproduce results are openly available on GitHub at https://github.com/erisemberg/peanut-allergy

## ACKNOWLEDGEMENTS

We thank UNC’s Systems Genetics Core Facility for maintenance and distribution of the Collaborative Cross mice. We thank the UNC Flow Cytometry Core Facility *(RRID:SCR_019170, supported in part by P30 CA016086 Cancer Center Support Grant to UNC’s Lineberger Comprehensive Cancer Center)* for access to their cytometers and advice in panel building.

