## Supplemental Figures for "A mutation in *Themis* contributes to peanut-induced oral anaphylaxis in CC027 mice"

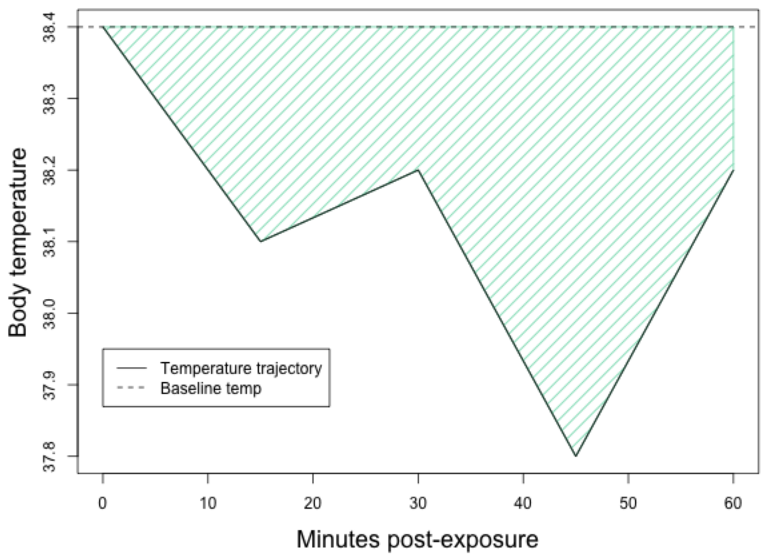


**Supplemental Figure 1. Illustration of area above the curve summary statistic**. Area above the curve (AAC) was calculated for each mouse by calculating the area between the horizontal line formed by the mouse's baseline temperature and the line formed by the temperature trajectory. It is possible to have a negative AAC if the temperature increases above baseline.

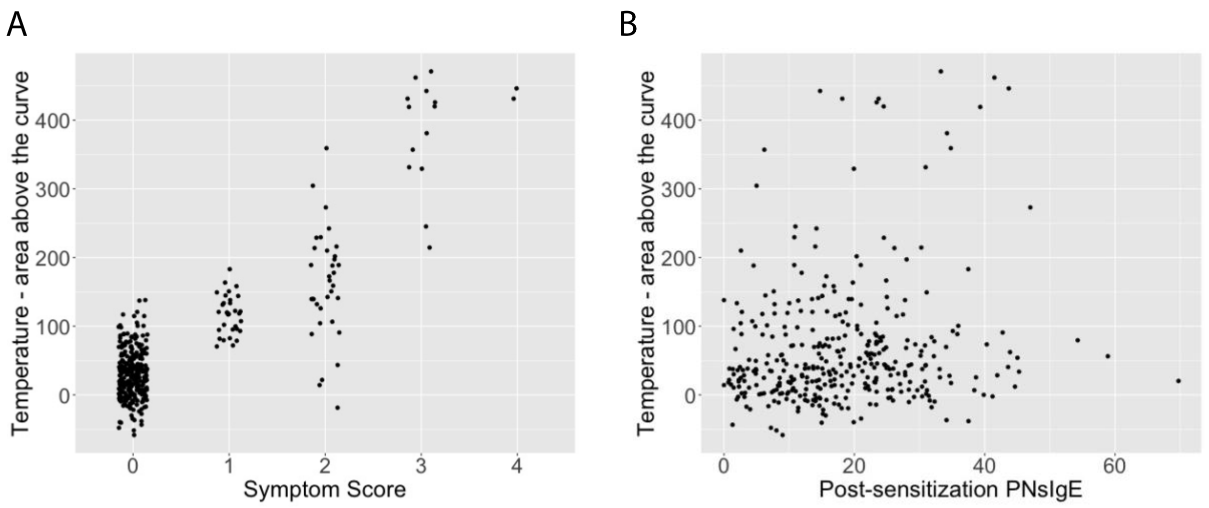


**Supplemental Figure 2. Correlation between measured phenotypes.** A) Symptom score and AAC are significantly associated ($p < 2x10^{-16}$, ANOVA). B) Post-sensitization peanut-specific IgE (PNsIgE) is weakly but significantly correlated with AAC ($R^2 = 0.16$, $p = 0.002$, Pearson's correlation test).

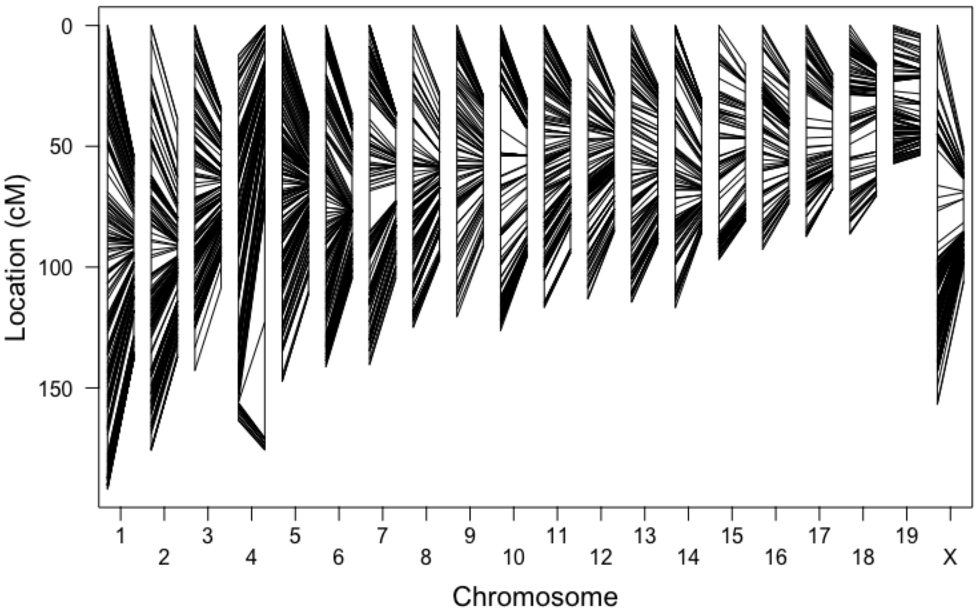


**Supplemental Figure 3. Genetic maps**. Observed (left) and estimated (right) genetic map for each chromosome, showing sufficient genome-wide marker density to perform QTL mapping. Estimated genetic map is produced with a hidden Markov model using the Lander-Green algorithm. Median physical distance between adjacent markers is 450 kilobases (kb) and the largest gap is 18 megabases (Mb) or approximately 9 centimorgans (cM) on chromosome 10.

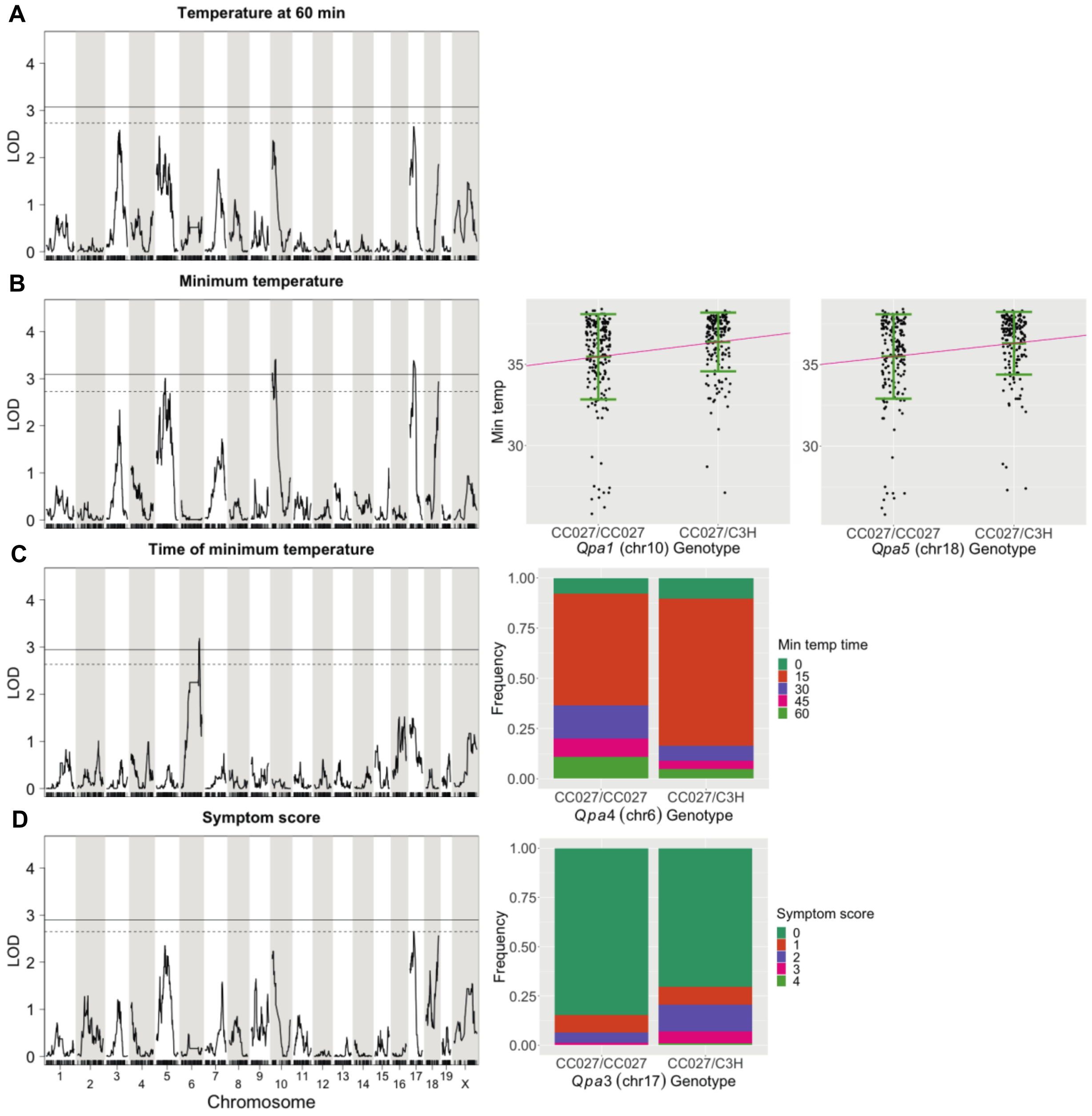


**Supplemental Figure 4. Additional temperature phenotypes**. Genome-wide scan for QTL associated with temperature phenotypes. Solid and dashed lines represent 5% and 10% significance thresholds, respectively. The right panel shows phenotype-by-genotype plots showing relationship between the genotype at each QTL and its associated phenotype. A) Temperature at 60 minutes. B) Minimum temperature. C) Time of minimum temperature. D) Symptom score.

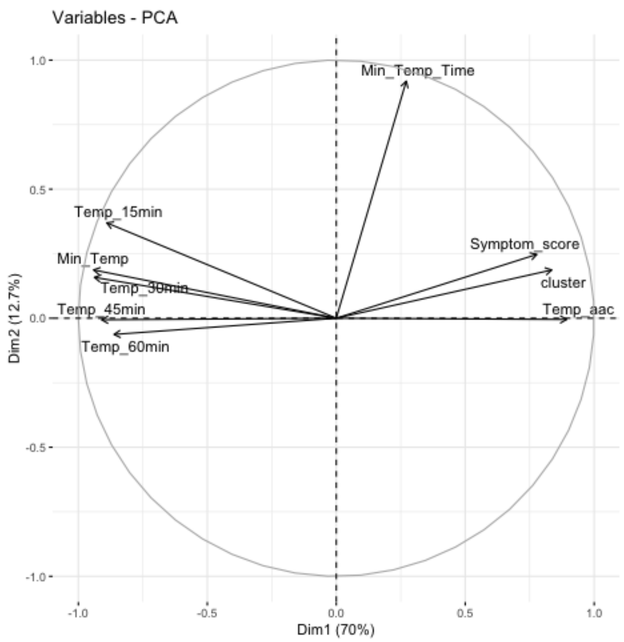


**Supplemental Figure 5. Variable PCA shows that time of minimum temperature captures variation distinct from the other temperature-related phenotypes**. Principal components analysis was performed on centered and scaled temperature phenotypes. Time of minimum temperature (Min_Temp_Time) is captured by PC2 while the remaining phenotypes associate with PC1.

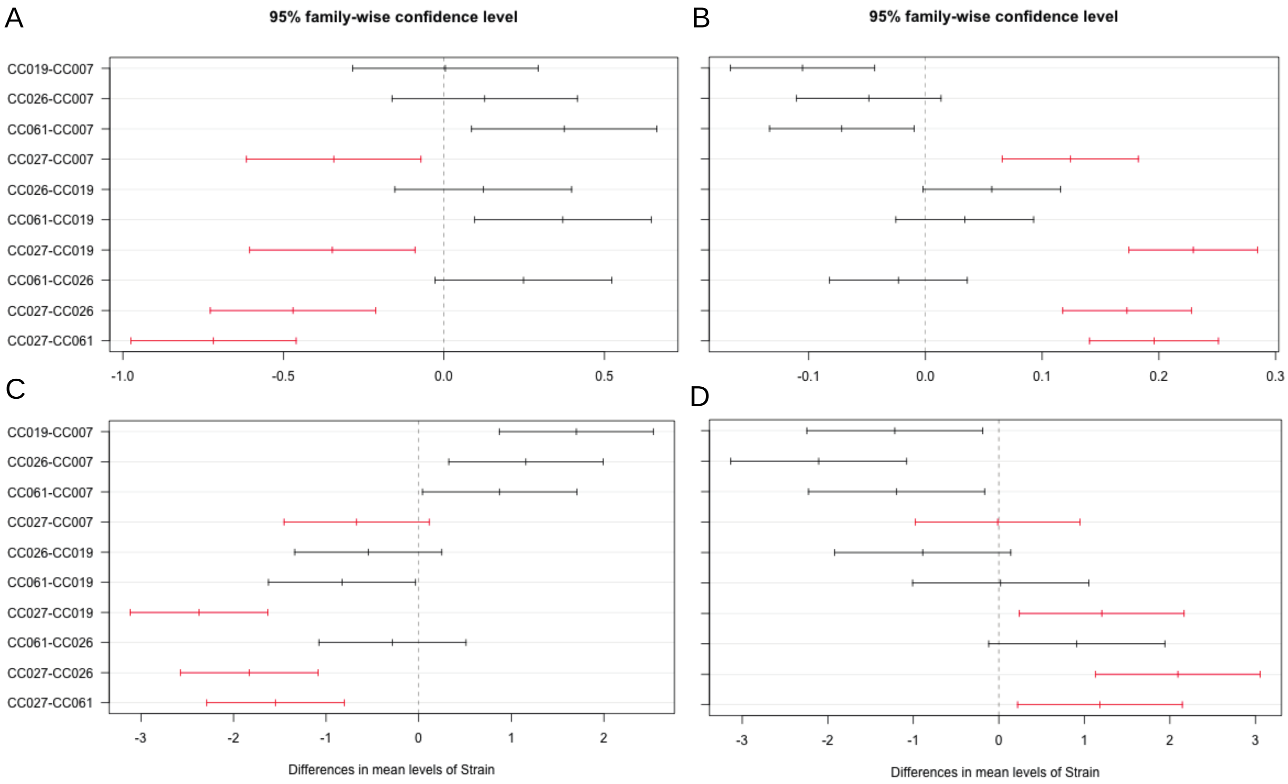


**Supplemental Figure 6. Tukey's Honest Significant Difference results**. Tukey's HSD test was performed as a more conservative test of mean differences which controls for type I error rate. Comparisons with CC027 are highlighted in red. A) Total regulatory T cells in the spleen (CC027 is different from all other strains). B) Proportion of double negative T cells in the spleen (CC027 is different from all other strains). C) Proportion of CD8+ T cells in the spleen (CC027 is different from all strains except CC007). D) Proportion of CD4+ T cells in the thymus (CC027 is different from all strains except CC007).
