## Supplemental Methods for "A mutation in *Themis* contributes to peanut-induced oral anaphylaxis in CC027 mice"

*Phenotype processing*

We calculated two summary statistics from raw temperature data. A statistic representing overall temperature trajectory for each animal was calculated by measuring the area between a horizontal line at baseline temperature and the line formed by the temperature trajectory (called area above the curve, or AAC, illustrated in **Supp. Fig. 1**). To identify groups of similar temperature trajectories, we performed complete linkage hierarchical clustering based on Euclidian distance between temperature measurements. This clustering was then binarized (cluster 1 vs clusters 2 and 3, representing non-reactors vs reactors) to improve power for genetic mapping. Raw temperature measurements, PNsIgE and AAC were normalized via truncated rank inverse normal transformation. Categorical (time of minimum temperature and symptom score) and binary (cluster) phenotypes were not transformed.

*Heritability Estimation*

The heritability, or the proportion of phenotypic variation explained by genetic variation, was estimated in two ways. First, the phenotypic variability between inbred parent strains was used to estimate narrow-sense (additive) heritability. For each phenotype, the following linear regression was performed:

$$y_i = \mu + \beta_{t0} x_{t0,i} + \beta_{\text{strain}} x_{\text{strain},i} + \varepsilon_i \,,$$

where $y_i$ is the phenotype of mouse $i$, $\mu$ is the intercept, $x_{t0,i}$ is the baseline temperature for mouse $i$, $\beta_{t0}$ is the effect of baseline temperature, $x_{\text{strain}}$ is the strain (CC027 or C3H) for mouse $i$ and $\beta_{\text{strain}}$ is the strain effect, and the residual $\varepsilon_i$ is normally distributed with variance $\sigma^2$. This model was used to calculate total sum of squares (TSS) and fitted sum of squares (FSS) for the strain term, and heritability was calculated as

$$h^2 = \frac{\text{FSS}_{\text{strain}}}{\text{TSS}} \,.$$

Next, the proportion of phenotypic variance explained by the additive effects at all genotyped SNPs (additive SNP heritability[38]) was calculated using the genetic and phenotypic similarity between backcross mice. The genomic relationship matrix, $\boldsymbol{G}$, was defined using GCTA-GRM as described in Yang et al (2011)[39]. The lme4qtl R package[40] was then used to fit the following model:

$$y_i = \mu + \beta_{t0} x_{t0,i} + g_i + \varepsilon_i \,,$$

where $g_i$, the polygenic effect for mouse $i$, follows a multivariate normal distribution with variance-covariance matrix $\boldsymbol{G}\sigma_g^2$. Additive heritability was then estimated as the intraclass correlation coefficient

$$h^2 = \frac{\sigma_g^2}{\sigma_g^2 + \sigma^2} \,.$$

*Quantitative Trait Loci (QTL) Mapping*

We performed QTL mapping to identify quantitative trait loci associated with our phenotypes. For our normally distributed (non-categorical) phenotypes, we first regressed out the effects of cage and batch from each phenotype using a random effects linear model with the phenotype as the outcome and cage and batch as random effects. That is, we fitted the model

$$\boldsymbol{y_{ijk}} = \boldsymbol{\mu} + \boldsymbol{b_j} + \boldsymbol{c_{jk}} + \boldsymbol{\varepsilon_{ijk}} \,, \tag{1}$$

where $y_{ijk}$ is the phenotype for mouse $i$ in batch $j$ and cage $k$, $\mu$ is the intercept, $b_j \sim N(0, \sigma_b^2)$ are the batch effects, $c_{jk} \sim N(0, \sigma_c^2)$ are the effects of cage (nested within batch), and $\varepsilon_{ijk} \sim N(0, \sigma^2)$ is the residual. The residuals estimated from this model were then used as the (batch- and cage-) corrected phenotype. For each corrected phenotype, we performed QTL mapping using Haley-Knott regression[41] as implemented in R/qtl[42], fitting at each genotyped marker a linear model that includes an effect of the marker genotype dosage. For temperature phenotypes, baseline temperature was also included as a fixed effect covariate, ie,

$$y_i = \mu + \beta_{\text{t0}} x_{\text{t0},i} + \beta_{\text{geno,j}} x_{\text{geno},ij} + \varepsilon_i$$

where $x_{\text{geno},ij}$ is the genotype dosage for mouse $i$ at marker $j$ and $\beta_{\text{geno,j}}$ is the effect of marker $j$. QTL mapping was performed on categorical phenotypes (time of minimum temperature, symptom score) using a nonparametric model and on binary phenotypes (binarized cluster) using a binary model as implemented in R/qtl[42].

Significance thresholds were determined by permutation test: in each of 1000 permutations, genotypes were shuffled with respect to phenotypes, breaking the relationship between the two and providing a null model appropriate for our phenotypic, experimental, and genetic data distributions. A genome scan was performed on this shuffled data and the highest genome-wide maximum LOD score was recorded. The distribution of 1000 maximum LOD scores was then used to fit a Generalized Extreme Value (GEV) distribution[43,44] using the extRemes R package[45]. This fitted GEV distribution was then used to calculate 5% and 10% significance thresholds for each genome scan and genome-wide adjusted p-values for each QTL. Since we only have female backcross mice, and only mice produced from female $F_1$ parents can be used for QTL mapping on the X chromosome, X chromosome-specific permutation tests are not necessary. Bayesian credible intervals were calculated for QTLs as initially described by Sen & Churchill (2001)[46] and as implemented in R/qtl[42].

*Phenotypic Variation Explained by QTL*

The proportion of phenotypic variation explained by a QTL (also referred to as the heritability due to the QTL) was calculated as follows for each phenotype associated with a QTL:

$$h^2_{\text{QTL}} = \frac{\text{var}\{\text{E}(y|g)\}}{\text{var}(y)} = \frac{a^2}{a^2 + 4\sigma^2},$$

where $y$ is the phenotype, $g$ is the QTL genotype, $a = \mu_{\text{AB}} - \mu_{\text{AA}}$ (the difference between the phenotypic means for heterozygous mice and homozygous mice), and $\sigma^2$ is the residual variance (phenotypic variance not explained by the QTL).

*Multiple QTL Analysis*

To determine the extent to which each QTL uniquely contributes to the associated phenotypes we performed variable selection via stepwise regression. The AAC statistic was chosen as a representative phenotype for the severity of allergic response. Principal components analysis (PCA) and factor analysis were used to verify that AAC was representative of all outcome phenotypes except time of minimum temperature. A linear model was then fitted with AAC as the outcome variable and every QTL (*Qpa1-6*) associated with anaphylaxis as predictor variables. We then performed stepwise regression using the stepAIC() function in the R package MASS[73], alternately removing and adding QTL to find a submodel that optimized the model fit as judged by the Bayes Information Criterion (BIC)[74].

*Analysis of Themis variant T-cell phenotypes*

To analyze the specific immune cell types where CC027 significantly differed from other strains, two analyses were performed after Z-score normalization: 1) a linear regression with Helmert contrasts, to determine whether the measurements for CC027 were different from the average of the other strains, and 2) Tukey's Honest Significant Difference[60], a conservative approach which tests all pairwise differences while controlling for the probability of making one or more Type I error in order to identify significant differences among sample means.

We performed PCA via singular value decomposition of the data matrix to assess the degree to which CC027 had a different T cell compartment overall relative to the other four strains analyzed. Prior to PCA, the data were Z-score normalized, and one mouse with missing data was removed (as data were not missing completely at random and so could not be reliably imputed).
